# Identification and full genome analysis of the first putative virus of sea buckthorn (*Hippophae rhamnoides* L.)

**DOI:** 10.1101/2022.07.27.501707

**Authors:** Ina Balke, Vilija Zeltina, Nikita Zrelovs, Ieva Kalnciema, Gunta Resevica, Rebeka Ludviga, Juris Jansons, Inga Moročko-Bičevska, Dalija Segliņa, Andris Zeltins

## Abstract

The agricultural importance of sea buckthorn (*Hippophae rhamnoides* L.) is rapidly increasing. Several bacterial and fungal pathogens infecting sea buckthorn have been identified and characterized; however, the viral pathogens are not yet known. In this study, we identified, isolated, and sequenced a virus from a wild plantation of sea buckthorn for the first time. Sequence analysis of the obtained viral genome revealed high similarity with sequences of several viruses belonging to the genus *Marafivirus*, especially olive latent virus 3 (OLV-3). The genome of the new virus is 6,989 nucleotides (nt) in length according to 5′ and 3′ rapid amplification of cDNA ends (RACE) (without polyA-tail), with 5′ and 3′ untranslated regions being 133 and 109 nt long, respectively. The viral genome encoded two open reading frames (ORFs). ORF1 encoded a polyprotein of 1,954 amino acids (aa) with the characteristic marafivirus non-structural protein domains—methyltransferase, Salyut domain, papain-like cysteine protease, helicase, and RNA-dependent RNA polymerase. ORF1 was separated from ORF2 by a six nt and encoded the coat protein (CP). CP had typical signatures of minor (30.96 kDa) and major (21.18 kDa) forms. Both CP forms were cloned and expressed in a bacterial expression system, and only the major CP was able to self-assemble into 30 nm virus-like particles that resembled the native virus, thus demonstrating that minor CP is not essential for virion assembly. We suggest the newly discovered virus to be named as “Sea buckthorn marafivirus”, abbreviated as “SBuMV”.

**Author summary:** Sea buckthorn is an exceptionally valuable plant that is currently widely cultivated as multipurpose horticultural species for food, pharmacology, cosmetics, and landscape conservation. Diseases and pests directly affect the cultivation of SBT. To date, several pests and diseases, mainly fungal and bacterial, but no viral, sea buckthorn have been reported. Identification of new pathogens would assist in the development of control strategies, and quarantine purposes and ensure sustainable sea buckthorn cultivation. Here, for the first time, we present a virus putatively infecting sea buckthorn. We had characterized its full genome, cloned and expressed minor and major forms of coat protein either individually or co-expressed. We also showed that major coat protein-derived virus-like particles self- assembled directly in the bacterial cells, and the majority of the expressed CPs were soluble. Our study suggests that the minor CP is not essential for the assembly of seemingly structurally intact viral particles, meaning that it can have other functions.

## Introduction

Sea buckthorn (SBT, *Hippophae rhamnoides* L., genus *Hippophae* L., family *Elaeagnaceae*) is an exceptionally valuable plant that is currently domesticated and cultivated in orchards, particularly in Europe, Canada, and the USA [1]. SBT is a spinescent, deciduous, and dioecious berry-producing shrub or small tree [2]. In addition, it is a pioneer species that is highly adaptable to extreme climatic and soil conditions and is tolerant to drought and extreme temperatures ranging from −43 °C to 55 °C [3, 4]. It is an ideal plant for soil erosion control, land recovery, wildlife habitat enhancement, and farmstead protection. SBT has attracted considerable attention from researchers, producers, and various industries because it has a high nutritional and medicinal value for humans [5]. SBT is not a native plant to Latvia, although fossil pollen records indicate its presence in postglacial raw soils in this region. The deliberate cultivation of SBT can be traced back to more than 100 years [6]. The first recorded trials with introduced SBT in Latvia were recorded in 1970s when the seedlings (‘Bogatirskaja’ and ‘Malutka’) originating from Russia (Kaliningrad region or Barnaul) were planted along the roadsides as windbreakers, as well as for recultivation of the exhausted dolomite, sand, and gravel pits [7, 8]. However, it was not until 1980 that SBT was grown as a plant of agricultural importance in Latvia [9].

To date, limited research related to disease and pest control has been reported in SBT due to the comparatively short time of domestication and cultivation in large orchards [10, 11]. Diseases and pests, which can alter almost the morphology and developmental stage of SBT, directly affect the cultivation of SBT. To date, several pests and diseases of SBT have been reported, and more are likely to be identified with the increase in the number of SBT plantations [11, 12]. The major fungal diseases reported in SBT include verticillium wilt, Fusarium wilt, stem canker, damping off, brown rot, scab, and dried-shrink disease. The common pathogenic fungi reported in SBT include species of the genera *Fusarium, Verticillium, Diaporthe, Hymenopleella, Alternaria, Pythium, Monilia, Valsa,* and *Stigmina* [11, 13]. Forty seven fungal species affecting SBT have been reported in Russia, with *Fusarium sporotrichiella* reported to cause the maximum damage [14]. In Finland, the genus *Stigmina* was reported to be the cause of stem canker, which can kill the entire shrub of susceptible cultivars [15]. The incidence of powdery mildew in SBT has been recorded in Himachal Pradesh [16]. Fungal endophytes (*Aspergillus niger*, *Mortierella minutissima,* and sterile mycelia of Basidiomycotina) and spores of four species of vesicular-arbuscular mycorrhiza (*Glomus albidum*, *G. fasciculatum*, *G. macrocarpum*, and *Gigaspora margariata*) have been isolated from different SBT plant parts and associated soil samples [17]. *Emericella quadrilineata* was isolated from the leaves of *H. salicifolia* in India [18]. Moreover, root rot caused by *Rhizoctonia solani* is a major problem in greenhouses in Uttarakhand [14]. In China, species of *Fusarium* and *Phomopsis* were found to be associated with dried-shrink disease in SBT [19]. Several fungi and bacteria (*Verticillium*, *Fusarium*, *Diaporthe*, *Eutypa, Hymenopleella,* and *Pseudomonas spp.*) were identified as possible causes of SBT decline in Latvia [20, 21]. Although many fungal species have been reported as pathogens of SBT, disease symptoms using bioassays and fulfilment of Koch’s postulates have been verified only for a few pathogens, including *V. dahliae*, *F. sporotrichioides, F. acuminatum, F. oxysporum, F. camptoceras*, *Stigmina* sp., and *Lepteutypa* sp. [11]. However, surprisingly, we did not find any reports of viruses infecting *H. rhamnoides*.

Since 2009, various RNA sequencing methods have been developed to identify phytoviruses, including sequencing of total RNA, ribosomal RNA-depleted total RNA, double-stranded RNA, virus-derived small interfering RNA, RNA from purified or partially purified viral particles, polyadenylated RNA, and RNA after subtractive hybridization with healthy plant RNA [22]. High throughput sequencing (HTS) technologies have quickly become a go-to “golden standard” method for novel virus identification and monitoring in various sample types. HTS eliminates the need for prior knowledge of the expected viral genomic sequences, thus providing a substantial advantage over traditional methods such as PCR amplification or microarray hybridization, which are dependent on target-specific primers [23]. HTS allows the identification of phytoviruses from various different material types at a potentially unlimited resolution. HTS has been successfully applied for such purposes in mixed infections [24], wastewaters [25], symptomless plants [26], herbarium [27], large field surveys [28], and human feces [29]. The plant virus genus *Marafivirus* is one of the many examples in which the introduction of HTS technology has resulted in the identification of multiple novel species. Consequently, the number of recognized *Marafivirus* species has increased from four (*Bermuda grass etched-line virus*, *Citrus sudden death-associated virus*, *Maize rayado fino virus*, and *Oat blue dwarf virus*)) in 2009 to 11 in 2020 (*Alfalfa virus F*, *Blackberry virus S*, *Grapevine asteroid mosaic associated virus*, *Grapevine Syrah virus 1* (also called *Grapevine virus Q*), *Nectarine marafivirus M*, *Olive latent virus 3*, and *Peach marafivirus D*) in addition to the previously mentioned species and not counting the tentative species [30].

The *Marafivirus* genus name was derived from the reference strain of the ***Ma***ize ***ra***yado ***fi***no virus [31]. Viruses belonging to the genus *Marafivirus* within the family *Tymoviridae* are small isometric plant viruses with a monopartite positive-sense single-stranded RNA genome (gRNA) varying from 6.3 kb to 7.1 kb in length (excluding the poly-A tail). gRNA has a 5′ cap, a 3′-part encoding a tRNA-like structure, and a poly-A tail [32]. Marafiviruses contain the “marafibox” sequence [CA(G/A)GGUGAAUUGCUUC] of 16 nt that is conserved and comparable to the “tymobox” (differs by three or four nucleotides), which has been shown to be a subgenomic promoter in tymoviruses [33]. Based on the predicted activity and location of the signature domains encoded within their genomes, marafiviruses are included in the alphavirus-like superfamily [34, 35]. However, viruses presently classified as marafiviruses exhibit some diversity in their genome architectures [36–40]. The very narrow host range of marafiviruses makes host susceptibility an important species demarcation criterion. The genomes of marafiviruses have a high cytidine (C) content (36–45%) and usually encode a single large precursor polyprotein containing methyltransferase (MT), Salyut domain (SD), papain- like cysteine protease (PRO), helicase (HEL), RNA-dependent RNA polymerase (RdRp), and coat protein (CP) domains. CP is commonly found in virions in two forms: major (small p21 kDa) and minor (large p23–25 kDa) [41, 42]. Interestingly, three CP fractions have been previously identified for oat blue dwarf virus (OBDV) and citrus sudden death-associated virus (CSDaV) after virus purification [38, 43]. Major and minor CPs were found in the virus particles at a molar ratio of approximately 3:1 [36, 44]. The minor CP is initially produced as a C-terminal fusion of the replication protein, whereas the major CP is produced from subgenomic RNA (sgRNA) [41]. The *in planta* experiments using the infectious cDNA of CSDaV revealed that major p21 is indeed a product of direct translation by leaky scanning from the second start codon in sgRNA. The minor CPs, p25 and p23, are produced by direct translation from the first start codon in sgRNA and by trans-proteolytic cleavage processing derived from the p25 precursor, but not as the fusion part of the polyprotein [43]. For OBDV CP, the major CP is translated directly from the sgRNA, while the minor CP is cleaved from both the polyprotein and a minor CP precursor translated from the sgRNA [45]. However, for Maize rayado fino virus (MRFV), minor and major CPs are largely translated from the sgRNA [46]. Unlike other marafiviruses, Alfalfa virus F (AVF) and putative marafivirus Medicago sativa marafivirus 1 (MsMV1) do not have a second initiation codon for the coding region of the major CP1 and only encode methionine (Met) for the minor CP [47, 48]. In this case, a possible strategy to produce the two CPs could be direct translation of the sgRNA for the minor CP and posttranslational cleavage of the larger precursor to produce a major CP [36]. The reason for the multiple expression strategies for minor CP is unclear. Edwards and Weiland [44] proposed that the cleavage of the replicase polyprotein does not produce stoichiometrically sufficient amounts of minor CP necessary for virion assembly. They also suggested an evolutionary transition toward CP production solely via sgRNAs and that the readthrough of a larger replicase polyprotein is vestigial. However, the production of a minor CP through a cleavage mechanism provides a regulatory feature with probable functional significance for both replication and encapsidation [44].

Marafiviruses are phloem-limited and, thus, are generally not sap transmissible. MRFV has been shown to be transmissible by vascular puncture [49, 50]. Marafiviruses are thought to be transmitted by leafhoppers in a persistent and propagative manner. Nevertheless, relatively few plant viruses naturally infect both insects and plant cells. Only the viral families *Rhabdoviridae*, *Reoviridae*, and *Bunyaviridae* and the genus *Marafivirus* are propagatively transmitted [51]. It was speculated that the complexity of CP expression in marafiviruses relative to that in tymoviruses might be related to the infection of both plant and insect hosts by marafiviruses [44]. Minor CP-depleted mutants of MRFV do not retain encapsidation and systemic infectivity, but they retain leafhopper transmissibility, indicating that the 37 amino acid (aa) N-terminal extension of minor CP is a leafhopper transmission signal sequence. Additionally, loss of major CP expression did not result in systemic infection [46].

In this study, we isolated an isometric virus approximately 30 nm in diameter from SBT plants obtained from a local wild plantation and sequenced its complete genome. HTS data analysis revealed that the isolated virus demonstrated genome and functional domain organization consistent with those of phytoviruses belonging to the genus *Marafivirus*. A close relationship with the recognized and tentative *Marafivirus* representatives was further supported by phylogenetic analyses of the genome and protein sequences of the novel virus. Based on the complete genome sequence obtained by combining RNA-seq, RT-PCR, Sanger sequencing, and 5′ and 3′ rapid amplification of cDNA ends (RACE), as well as the genome organization of the isolated virus, we propose that the new virus can be considered a novel viral species within the genus *Marafivirus* and should be named as Sea buckthorn marafivirus (SBuMV) isolate BU1. Additionally, cloning and expression of both *CP* gene product variants (minor and major) of SBuMV in *E. coli* expression system were investigated.

## Materials and Methods

### Plant source

Leaf material of SBT was collected in August, 2017 and August, 2021 from the same three individual SBT shrubs with leaves bearing necrotic spots. SBT source plants were sampled at a wild plantation grown since at least the 1950s at the Buļļupe river bank to control soil erosion (GPS location:57.018110N; 24.004251E).

### Virus and RNA purification

The viral fraction was separated from 30 g of leaf material of each SBT sample that was collected in August, 2017. Leaves were grinded in the electric mixer ETA 0010 (ETA, Prague, Czech Republic) in 100 ml of homogenization buffer [0.02 M HEPES pH 6.8, 0.2 M sucrose, 20 mM sodium azide (NaN_3_), 10 mM β-mercaptoethanol (β-ME), 1 mM PMSF, 2 mM EDTA]. The suspension was centrifuged in a low-speed centrifuge 5804R (Eppendorf, Hamburg, Germany) at 11,000 rpm (15,557 × *g*) and 4 °C for 15 min. The resulting supernatant was filtered using a filter paper. TX-100 was added to the clarified supernatant drop by drop on a magnetic stirrer at 4 °C until the 5% saturation was reached. The solution was then loaded on a 20% sucrose cushion in a 0.02 M HEPES buffer (1:1 volume ratio; pH 8.2) and sedimented by ultracentrifugation on Optima-XL (Backman Coulter, Brea, CA, USA) using Type-70Ti rotor (Backman Coulter, USA) at 50,000 rpm (183,960 × *g*) and 4 °C for 1.5 h. The obtained supernatant was discarded and the pellet was dissolved in 6 ml of buffer [0.1 M sodium borate (NaB) pH 6.8, 5 mM EDTA, overnight (ON)] at 4 °C. The solution was then clarified by centrifugation in a low-speed centrifuge 5804R (Eppendorf, Hamburg, Germany) at 10,000 rpm (12,857 × *g*) and room temperature for 5 min. Soluble fraction (6 ml) was further purified on a sucrose gradient containing layered sucrose fractions ranging from 10% to 40% (with intervals of 10%) in 0.1 M NaB (pH 8.2). Sucrose gradient centrifugation was performed as previously described [52]. After sucrose gradient centrifugation, 6.5 ml fractions from the bottom of the tubes were carefully collected. Sucrose fraction analysis was performed using sodium dodecyl sulphate–polyacrylamide gel (12.5%) electrophoresis (SDS-PAGE) followed by Coomassie G250 (Sigma-Aldrich, St. Louis, MO, USA) staining. Two of the fractions (40% and 30%) were dialyzed against 100 volumes of 0.05 M NaB (pH 8.2) ON on a magnetic stirrer. Dialyzed solution was loaded on a 20% sucrose cushion in 0.05 M NaB and 2 mM EDTA and sedimented by ultracentrifugation on Optima-XL (Backman Coulter, Brea, CA, USA) in Type-70Ti rotor at 50,000 rpm (183,960 × *g*) and 4 °C for 4 h. The supernatant was removed and the pellet was dissolved in 0.3 ml solution containing 0.05 M NaB and 2 mM EDTA. The contents of the dissolved pellet solution were further analyzed using transmission electron microscopy (TEM). RNA was extracted from 100 µl of purified virus sample using the innuPREP Virus DNA/RNA Kit (Analytik Jena, Jena, Germany) according to the manufacturer’s protocol. The purified RNA was then analyzed using a NanoDrop-1000 spectrophotometer (Thermo Fisher Scientific, Waltham, MA, USA) to evaluate the average RNA concentration and assess the quality of the isolated RNA at 260/280 nm. Next, the purified RNA specimen was analyzed using an Agilent 2100 bioanalyzer (Agilent Technologies, Santa Clara, CA, USA) with an Agilent RNA 6000 Pico kit (Agilent Technologies, Santa Clara, CA, USA) to evaluate the fragmentation level of extracted RNA and length distribution of the obtained RNA fragments. Precise total RNA concentration was determined using a Qubit 2.0 (Thermo Fisher Scientific, Waltham, MA, USA) with a Qubit RNA high sensitivity (HS) assay kit (Thermo Fisher Scientific, Waltham, MA, USA).

For leaf samples collected in August, 2021, total RNA was isolated using the RNeasy Plant Mini Kit (Qiagen, Hilden, Germany) according to the manufacturer’s protocol. Purified RNA was then analyzed using a NanoDrop-1000 spectrophotometer (Thermo Fisher Scientific, Waltham, MA, USA) to assess the quality of isolated RNA at 260/280 nm. RNA concentration was determined using a Qubit RNA broad range assay Kit (Thermo Fisher Scientific, Waltham, MA, USA) on a Qubit 2.0.

### Transmission electron microscopy (TEM)

Purified virus or VLP samples were adsorbed onto carbon formvar-coated grids (300 mesh Copper/Palladium 3.05 mm; Laborimpex, Forest, Belgium) or carbon formvar-coated grids (hexagonal 200 mesh nickel 3.05 mm; Laborimpex, Forest, Belgium) for 1 min. The grids were then washed three times with 1 mM EDTA solution by brief immersion in the solution. The droplet was removed using a filter paper. Washed grids with the adsorbed samples were then negatively stained with a 0.5% uranyl acetate aqueous solution for 30 s. The stained droplet was removed using a filter paper. The samples were allowed to dry on filter paper in a Petri dish for 1 h before performing TEM. Grids were examined using a JEM-1230 electron microscope (JEOL, Tokyo, Japan) at an accelerating voltage of 100 kV.

### Sea buckthorn marafivirus (SBuMV) gRNA RNA-seq library preparation for HTS on the MGI platform

The HTS library was generated for the viral RNA extracted from the pooled plant samples collected in August, 2017 using the MGIEasy RNA Directional Library Prep Set for 16 reactions (MGI, Shenzhen, China) according to the manufacturer’s protocol for pair-end reads of 150 bp. Libraries were verified on an Agilent 2100 bioanalyzer with a High Sensitivity DNA Kit (Agilent Technologies, Santa Clara, CA, USA) for the average fragment length calculation and evaluation of quality and on Qubit 2.0, with Qubit 1X dsDNA HS Assay Kits (Thermo Fisher Scientific, Waltham, MA, USA) for the measurement of library fragment concentration. Library pooling, circularization, and cleaning were performed according to protocol guidelines. The final concentration of the resultant library was measured with a Qubit ssDNA Assay kit (Thermo Fisher Scientific, Waltham, MA, USA), and HTS was performed on a flow cell PE150 (MGI, Shenzhen, China) using a DNBSEQ-G400 (MGI, Shenzhen, China) system.

### SBuMV genome *de novo* assembly and preliminary annotation

Demultiplexed read data were inspected using FastQC v0.11.5 [53]. Transcriptome *de novo* assembly was performed using rnaSPAdes v3.13.1 [54]. Initial functional annotation of transcripts was performed using the ORF finder [55] and conserved domain search [56] of NCBI within the putative products of the predicted ORFs. Based on the conserved domain content of their putative products, transcripts of presumably viral origin were subjected to a BLASTn [57] search against the non-redundant nucleotide database (nr/nt) restricted to sequences of viral origin (taxid:10239) to gain an overview of their placement within the known virus diversity. Pairwise alignment of viral transcripts to those of the respective highest-scoring BLASTn hits was performed using EMBOSS Needle [58].

### SBuMV gRNA 5′ and 3′ RACE and gRNA RT-PCR fragment verification by Sangers sequencing

Prior to starting the 5′/3′-end RACE, the isolated viral RNA was tested by one-step RT-PCR using the Verso 1-Step RT-PCR Kit (Hot Start) (Thermo Fisher Scientific, Waltham, MA, USA) and nested primers—MarBox1F and MarBox1R. The corresponding PCR product (428 bp) was extracted from the gel after analysis in 0.8% native agarose gel (NAG) using the GeneJET Gel Extraction Kit (Thermo Fisher Scientific, Waltham, MA, USA) and cloned into the linearized vector pTZ-57 using InsTAclon PCR Cloning Kit (Thermo Fisher Scientific, Waltham, MA, USA). Conventional sequencing transformations into XL1-Blue Super competent cells (Agilent Technologies, Santa Clara, CA, USA) were used for all ligations. PCR fragment-containing clones were selected by digestion analysis using NcoI and XhoI restriction enzymes (Thermo Fisher Scientific, Waltham, MA, USA). Three positive clones were sequenced by the Sanger sequencing method using the ABI PRISM BigDye Terminator v3.1 Ready Reaction Cycle Sequencing Kit (Thermo Fisher Scientific, Waltham, MA, USA) and ABI PRISM 3130xl sequencer (Thermo Fisher Scientific, Waltham, MA, USA) with the corresponding primers M13seq-F and M13seq-R. Sequence assemblies were prepared using SeqMan software. To determine the complete genome sequence of SBuMV up to the last base, the 5′ and 3′’ untranslated regions (UTRs) of the virus were determined using a SMARTer®RACE 5′/3′ Kit (Takara Bio, USA). Viral RNA isolated in August, 2017 was used for UTR elucidation and resequencing. First-strand cDNA was amplified according to the manufacturer’s protocol. The 5′ (Fig 1C, 1) and 3′ (Fig 1C, 7) ends were amplified with the genome-specific primers MetPro-seq2-R and MarRdRp-seq2-F, respectively, and 10xUPS (SMARTer®RACE 5′/3′ Kit). An additional amplification step was employed because PCR products could not be initially detected in NAG. The second PCR was performed with Green Phusion polymerase (GFpol, Thermo Fisher Scientific, Waltham, MA, USA) according to the manufacturer’s protocol using the same template as used for the first PCR, the MarCP- 2F (3′ end RACE), and short UPS (SMARTer®RACE 5′/3′ Kit) or the first-strand cDNA, primer MetPro-seq1-R (5′ end RACE), and short UPS. PCR reactions were analyzed in 0.8% NAG, and the corresponding PCR products were extracted from the gel; adenine overlaps were added using *Taq* polymerase (Thermo Fisher Scientific, Waltham, MA, USA) for cloning into the linearized vector pTZ- 57 or directly after the purification into the linearized vector pJET1.2 using the CloneJET PCR Cloning Kit (Thermo Fisher Scientific, Waltham, MA, USA). Insert-containing clones were selected by restriction testing using EcoRI and HindIII restriction enzymes. At least three clones with the characteristic restriction pattern and direct PCR products were sequenced using the primers M13seq-F and M13seq-R for pTZ57 and pJET1.2-F and pJET1.2-R for pJET1.2. The generated sequences were aligned to the *de novo* assembled sequences from HTS data. The corresponding replication-associated polyprotein (RP) domains and CP were further amplified and verified by Sanger sequencing (Fig 1). The 5′ UTR-Pro fragment (Fig 1C, 2; 2023 bp) was amplified using 5′ RACE cDNA with GFpol in GC buffer and the primers Mar-5UTR-RACE-F and MetPro-seq2-R. The PRO domain (Fig 1C, 3; 1775 bp) was amplified using 5′ RACE cDNA with GFpol in GC buffer and the primers Mar-Pro2-F and Mar-Pro2-R. The Hel-RdRp domain (Fig 1C, 4; 2861 bp) was amplified using 3′ RACE cDNA with GFpol and primers Mar-Hel-F and Mar-RdRp-R, and the CP-3′ UTR fragment (Fig 1C, 6; 975 bp) was amplified using 3′ RACE cDNA with GFpol in GC buffer and primers Mar-CPL-F and Mar- 3UTR-RACE-R. The RdRp-C-term-CP fragment (Fig 1C, 5; 1066 bp) was amplified using 3′ RACE cDNA with GFpol in GC buffer and the primers SB-RdRp-CP-seq-F and MarCP2R. All corresponding PCR products were extracted from NAG and cloned into pTZ-57 or pJET1.2 cloning vectors. Clone selection and sequencing were performed as described above. MT pTZ-57 clones were sequenced with M13seq-F and M13seq-R; PRO pTZ-57 clones were sequenced with M13seq-F, M13seq-R, and internal primers—MetPro-seq2-F and MetPro-seq2-R—employing 5% dimethyl sulfoxide (DMSO) and a heating step (5 min at 95 °C) before the main sequencing program [59]. HEL-RdRp pJET1.2 clones were sequenced using the pJET1.2-F, pJET1.2-R, and internal primers—MarHel-seq3F and MarHel-seq4F—with a prior heating step and 5% DMSO, MarRdRp-seq2-F, MarRdRp-seq2-R, MarRdRp-seq3-R, and MarRdRp-seq5-R. CP-3′-UTR pJET1.2 clones were sequenced using pJET1.2- F and pJET1.2-R. The 5′-UTR-Pro pJET1.2 clones were sequenced using pJET1.2-F, pJET1.2-R, and the internal primers—MetPro-seq1-F and MetPro-seq1-R. RdRp-C-term-CP pJET1.2 clones were sequenced using pJET1.2-F and pJET1.2-R. All the primers used are listed in S1 Table.

**Fig 1.**
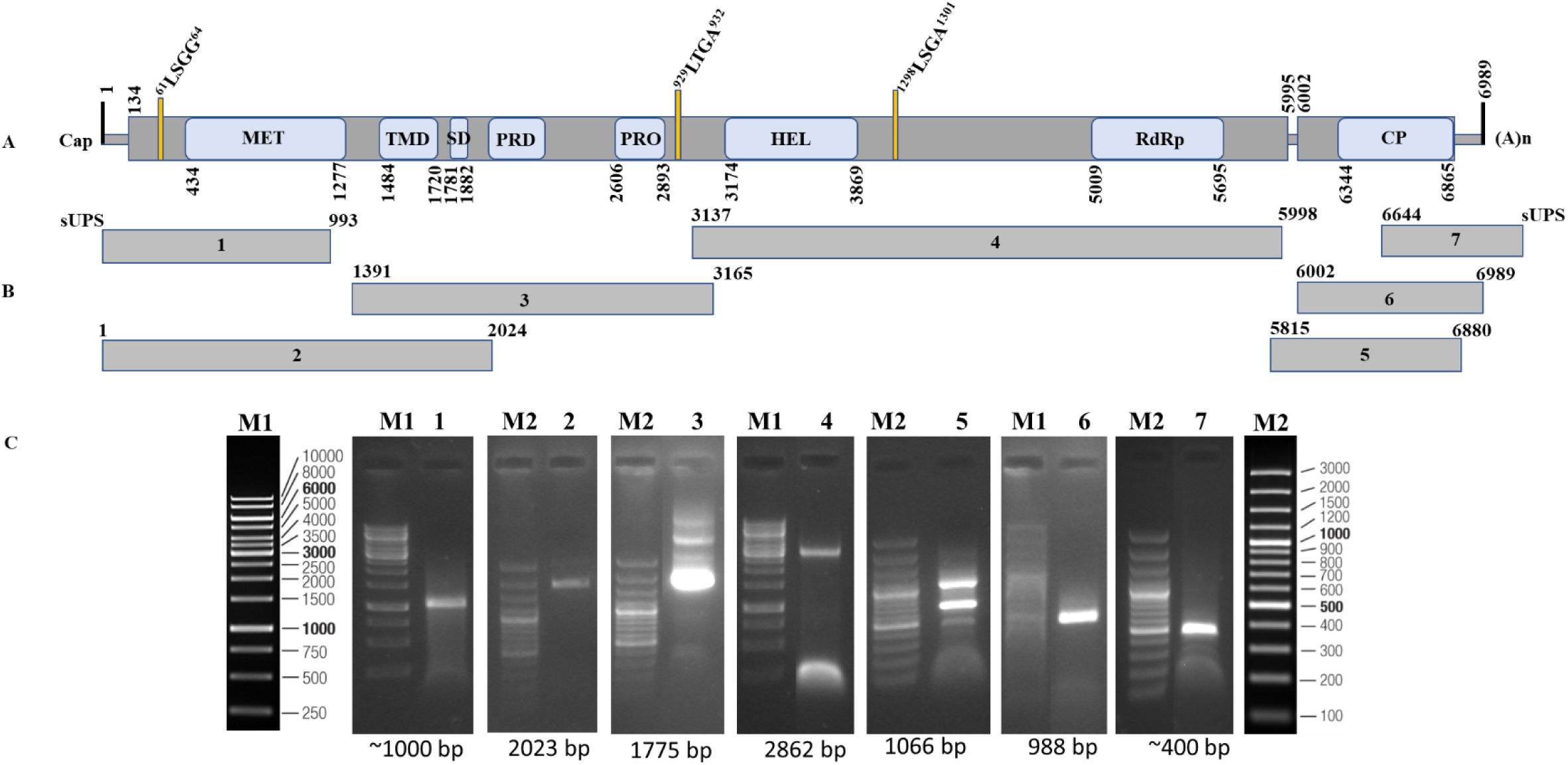
Schematic diagram of the SBuMV genomic sequence verification. MET – methyltransferase; TMD – transmembrane domain; SD - Salyut domain; PRD – proline rich domain; PRO - papain-like cysteine protease; HEL - helicase, RdRp - RNA-dependent RNA polymerase; CP - coat protein (see S1 Table for additional details).

### *Tymoviridae* sequence dataset acquisition

For reconstruction of SBuMV evolutionary relationships with other *Tymoviridae* viruses, a dataset (S2 Table) comprising public biological sequence repository accessions was additively built as follows: 1) exemplar isolates of species belonging to the family *Tymoviridae* from the Virus Metadata Repository number 18 of ICTV (October 19, 2021; MSL36); 2) *Tymoviridae* entries from RefSeq [60] (including those that do not yet have a standing in the official virus taxonomy as per ICTV Master Species List 2021.v1 (April 1, 2022); 3) *Tymoviridae* entries longer than 6000 bases representing presumably complete or near-complete genomes from the NCBI nucleotide database [61]. Respective nucleotide sequences were retrieved along with their taxonomy and the replicase/polyprotein and/or capsid protein sequences they encode (S2 Table). As some recognized and tentative *Marafiviruses* are known to have their CP encoded within the polyprotein gene, polyprotein sequences from entries that did not have an individual ORF encoding for CP were subjected to conserved domain prediction using batch CD-Search under default settings [62]. Entries with a discernible CP domain at the C-terminus of the polyprotein sequence were then partitioned by moving the last 300 aa of the polyproteins into the presumable CP aa sequence entries. Thus, for entries with CP encoded within the C-terminus of replicase polyprotein, truncated polyprotein aa sequence (without the last 300 aa; qualifier “_woCP” added to the original protein sequence accession in the label) was used for replicase phylogenies, and C-terminal sequences of 300 aa were used for CP phylogenies (qualifier “_300last” added to the original protein sequence accession in the label). MAFFT v7.453 [63] was used to generate multiple sequence alignments (MSAs) of the 1) complete or near-complete genome (hereafter referred to as genome MSA and, accordingly, genome tree), 2) polyprotein (without CP domain) aa, and 3) CP (including polyprotein-derived CP sequences) aa in automatic mode. Thereafter, each MSA was used to generate the respective maximum-likelihood and neighbor-joining trees.

### Evolutionary relationship analysis of the recognized and tentative *Tymoviridae* representatives

Maximum likelihood (ML) trees were constructed using IQ-TREE v. 2.0.3 [64], and ModelFinder [65] was used for the best substitution model selection according to the Bayesian Information Criterion, allowing for polytomies and using 1000 ultrafast bootstrap (UFBoot; [66]) replicates to determine the branch supports. Neighbor-joining (NJ) trees [67] were constructed using MEGA v 7.0.26 [68], eliminating all MSA positions with less than 90% site coverage, assuming uniform substitution rates, and determining branch supports using 1000 bootstrap [69] replicates. The trees were rooted using the respective outgroup (Botrytis virus F) sequences and visualized using FigTree v 1.4.4 [70]. Respective ML and NJ trees were then placed side-by-side and annotated using Inkscape v 1.0.1 [71], with distal nodes of the well-supported branches (≥95% UFboot for ML trees and ≥80% bootstrap for NJ trees) being colored in green. The technical parameters of the MSAs and inferred trees are shown in S3 Table.

### Expression, purification, and analysis of SBuMV CP cloned into the bacterial expression vector

Minor (p31) and major (p21.2) *CP* genes, which were obtained from the RNA extracted from the samples collected in August, 2017, were amplified by RT-PCR using the Verso 1-Step RT-PCR Kit with Thermo-Start Taq (Hot Start; Thermo Fisher Scientific, Waltham, MA, USA) using primers Mar- CPL-F and MarCP2R (876 bp) for the *p31* variant, and MarCP1F and MarCP2R (602 bp) for the *p21.2* variant. PCR products were cloned into the pTZ-57R/T vector (Thermo Fisher Scientific, Waltham, MA, USA). Clones containing *CP* inserts were identified by analyzing the pattern generated upon digestion with restriction enzymes NcoI and HindIII and confirmed by Sanger sequencing using M13seq-F and M13seq-R primers. Plasmid-harboring clones containing a *p31* or *p21.2* insert were digested with NcoI and HindIII, and DNA fragments were purified and cloned into the NcoI and HindIII sites of *E. coli* expression vector pRSFDuet1 (Novagen, Bad Soden, Germany), resulting in the plasmids pRSFDu-p31 or pRSFDu-21.2, respectively. For p31 and p21.2 co-expression, *p21.2* was amplified with the MARCP-short-Nde-F and MARCP-short-Xho-R primers. The PCR products were cloned into the pTZ-57R/T vector. Clones containing the *p21.2* insert were identified by analyzing the pattern generated upon digestion with restriction enzymes NdeI and XhoI and confirmed by Sanger-based sequencing using M13seq-F and M13seq-R primers. A plasmid-harboring clone containing a *p21.2* insert was digested with NdeI and XhoI, and DNA fragments were purified and cloned into the NdeI and XhoI sites of the *E. coli* expression vector pRSFDu-p31, resulting in the plasmid pRSFDu- p31-p21.2. Plasmid clones without sequence ambiguities were introduced into the C2566 *E. coli* expression strain (New England Biolabs, Ipswich, MA, USA). Cell cultivation and expression of CPs as well as cell disruption conditions were the same as previously described for cocksfoot mottle virus and rice yellow mottle virus [52]. VLPs were purified and analyzed in a similar manner to native virions, except that 1x PBS, 5 mM β-ME, and 0.5% TX-100 were used (see Virus and RNA purification).

### SBuMV detection by RT-PCR

The total RNA isolated from SBT samples collected in August, 2021 was used for SBuMV detection. One-step RT-PCR using a SuperScript™ III One-Step RT-PCR System with Platinum™ Taq DNA Polymerase (Thermo Fisher Scientific, Waltham, MA, USA) was performed using a combination of primers [Mar-CPL-F and MarCP2R (876 bp)] designed specifically for minor CP amplification.

## Results

### Virus purification from SBT leaf samples

The leaf material collected from the SBT shrubs was used for virus purification. Homogenization buffer (100 mL) used for leaf homogenization was concentrated 333 times using several purification steps, including sucrose cushions and sucrose gradients. TEM analysis revealed the presence of icosahedral viral particles in all the samples (Fig 2A).

**Fig 2.**
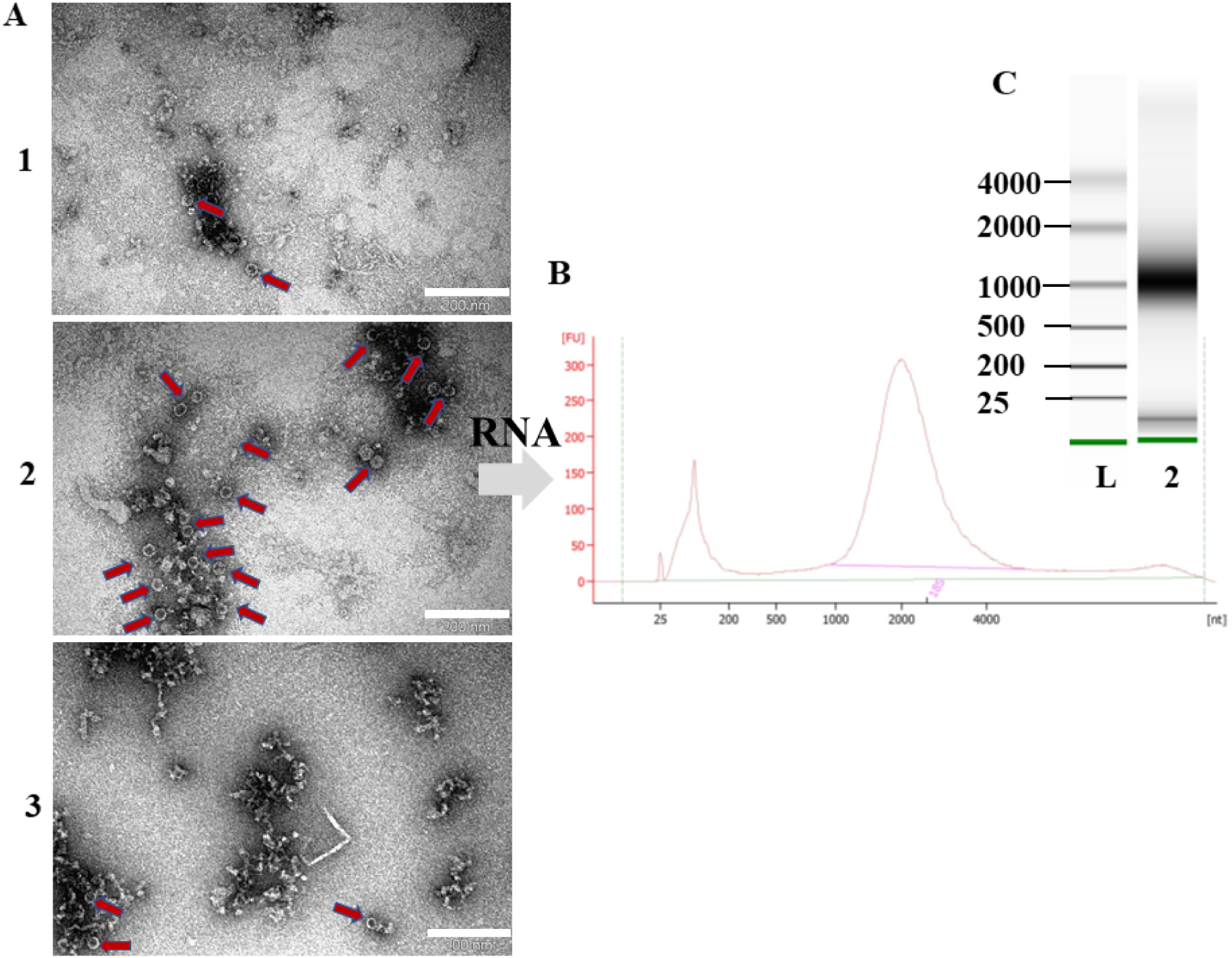
Purification of viruses from SBT and analysis of isolated RNA. A - TEM analysis of purified and concentrated SBT samples, scale 200 nm; B - chromatogram of isolated RNA from corresponding Bioanalyzer assays; C - electropherogram of isolated RNA from corresponding Bioanalyzer assays; 1– 3 - SBT samples; L – RNA ladder (Agilent Technologies, Santa Clara, CA, USA).

### SBuMV genome assembly

Demultiplexed HTS read data inspection using FastQC v0.11.5 [53] revealed that the sequencing run yielded 13,184,928 read (of up to 150 bp in length) pairs that were considered to be of sufficient quality for downstream processing. Transcriptome *de novo* assembly was performed using rnaSPAdes v3.13.1 [54], generating 147,111 transcripts of up to 47,770 bp long. Four of the *de novo* assembled transcripts (6,959–6,895 bp; 123–178 k-mer coverage) were identified to represent the near-complete genome variants of a putative novel *Tymoviridae* representative based on the characteristic presence of the conserved domain in their aa sequence of the putative ORF product. As these transcripts differed by multiple indels and single nucleotide polymorphisms, indicating extensive quasispecies within the host, oligonucleotide primers for both genome termini RACE and validation of the genome were designed based on the sequence of one of these near-complete genome transcripts that were selected randomly. After Sanger-based sequencing and mapping of the obtained reads onto the selected transcript that validated the sequence and allowed to extend the transcript, the genome of this novel putative *Tymoviridae* family representative was determined to constitute 7,020 bases (including 3’ poly-A tail) with a 53.8% GC content that had an average HTS read coverage of 819x. ORF prediction and functional annotation were performed using ORF finder tool [55] and conserved domain search of NCBI [56]. BLASTn [57] search against the nr/nt database restricted to the sequences of viral origin (taxid:10239) showed the highest total-scoring hit to the complete genome (7,148 bases) of OLV-3 isolate CN/1/1 (Accession: FJ444852.2), which had a query coverage and percent identity (e-value 1e- 150) of 60% and 68.81%, respectively. Lower-scoring hits were found to isolates of other *Marafiviruses* [e.g., grapevine asteroid mosaic-associated virus (GAMaV), OBDV, Nectarine marafivirus M (NeVM), and CSDaV]. Pairwise alignment of this novel virus genome to that of OLV- 3 using EMBOSS Needle [58] showed a length of 8,065 bases, 4630/8065 (57.41%) identities and 1962/8065 (24.33%) gaps. The full sequence of SBuMV was submitted to GenBank on March 31, 2022, and was assigned the accession number ON149451.

### SBuMV evolutionary relationships with other viruses

In all three of the tree pairs (genome sequence, CP, and replicase aa sequence), SBuMV shared a well-supported most recent common ancestor (MRCA) with OLV-3 isolate CN1/1 and was grouped in a major *Marafivirus* clade comprising all the taxonomically recognized marafiviruses, with the exception of AVF and some tentative marafiviruses (e.g., MsMV1, Davidia involucrata marafivirus 1 (DiMV1), and Glehnia littoralis marafivirus; Fig 3, S1 and S2 Figs).

**Fig 3.**
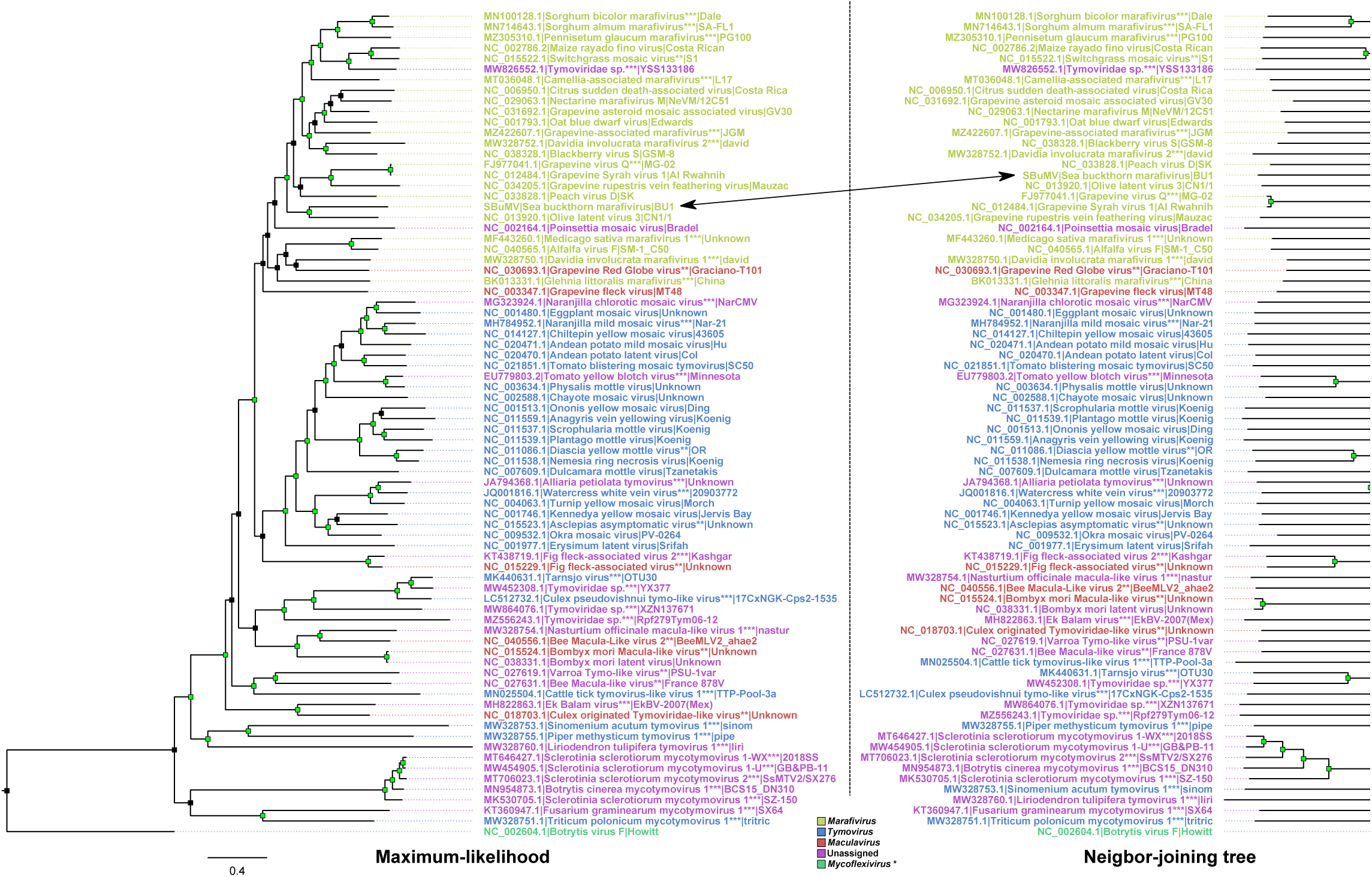
Maximum-likelihood and neighbor-joining trees of *Tymoviridae* complete or near- complete genome sequence. Trees are drawn to scale, with branch lengths in units of nucleotide substitutions per site. Tree tip labels are in the form of sequence accession number|virus name|strain and are colored based on the genera to which the virus belongs according to the legend. Two asterisks (**) after the virus name indicate that the virus originated from RefSeq and did not yet have a standing in the official virus taxonomy but was included in the analysis based on taxonomy associated with a RefSeq entry. Three asterisks (***) after the virus name indicate that the sequence originated from the Nuccore database and did not yet have a standing in the official virus taxonomy but was included in the analysis based on taxonomy associated with a complete or near-complete genome GenBank entry that was longer than 6000 bases in length. Botrytis virus F, which is a member of family *Gammaflexiviridae*, genus *Mycoflexivirus*, serves as an outgroup at which the trees are rooted. Black arrow connects Sea buckthorn marafivirus (SBuMV) leaf in both the trees.

This aforementioned major clade containing all the officially recognized marafiviruses (except for the AVF) was well supported (≥80% bootstrap support for the NJ trees; ≥95% UFBoot support for ML trees) in all the trees, except for the CP ML tree, and also included sequences from an unassigned *Tymoviridae* representative (isolate YSS133186) that had a well-supported common ancestry with MRFV isolate Costa Rican and some of the tentative marafiviruses. This suggests the possibility of classification of *Tymoviridae* isolate YSS133186 as a representative of *Marafivirus* genus. Interestingly, the AVF isolate SM-1_C50, which is recognized as a marafivirus, was not grouped in the large *Marafivirus* clade with sufficient support in either of the six trees, along with some other yet unrecognized tentative marafiviruses (based on the respective sequence submission-associated taxonomy; e.g., MsMV1, DiMV1 isolate david, and Glehnia littoralis marafivirus isolate China) that tended to cluster together, although they did not have a well-supported MRCA. Additionally, all the ICTV-recognized *Tymovirus* representatives formed a well-supported clade in all the trees, except for the CP NJ tree, with two of the tentative *Tymovirus* representatives forming a distinct clade comprising Sinomenium acutum tymovirus 1 isolate sinom and Piper methysticum tymovirus 1 isolate pipe (only shown in the genome and polyprotein tree as no CP sequence was readily available). Most of the “mycotimoviruses” that were listed as tentative *Tymoviridae* representatives without a genus-level assignment based on the submitted sequence-associated taxonomy [72], also formed a well-supported distinct clade in genome and polyprotein trees and might represent a putative novel genus. In all trees, various arthropod-associated tentative *Tymoviridae* representatives formed several distinct well- supported clades comprising a small number of leaves, and the majority of the sequences representing these leaves did not have a genus assigned to them. However, some of them were provisionally assigned to *Maculavirus* (e.g., Macula-Like virus 2, Bombyx mori Macula-like virus, and Culex originated Tymoviridae-like virus) and *Tymovirus* (e.g., Tarnsjo virus isolate OTU30, Cattle tick tymovirus-like virus 1, and Culex pseudovishnui tymo-like virus) genera as per NCBI taxonomy [72]. Overall, the results of our phylogenetic analyses suggest that some of the *Tymoviridae*-related virus sequences, without standing in the official virus taxonomy, have accumulated in the public biological sequence repositories, which, we believe, requires the attention of the ICTV representatives to formalize their place within the scope of the official virus taxonomy, thus encouraging further *Tymoviridae*-like virus diversity studies and clarifying the uncertainty regarding the intrafamily relationships of tentative *Tymoviridae* representatives that have been recently revealed.

### SBuMV genome annotation, resequencing, and 5′ and 3′ end mapping with RACE

To complete the *de novo* assembled viral genome, the 5′ and 3′ terminal ends and internal genome fragments were resequenced and validated using Sanger-based sequencing. To sequence both flanking ends of the viral genome, we used the switching mechanism at 5’ end of RNA template (SMART) RACE method. The 5′ or 3′ ends were amplified with custom SBuMV nested primers designed using processed RNA-seq data (*de novo* assembled transcripts) and SMARTer® RACE 5′/3′ kit primers. Custom primers for nested SBuMV PCR fragment amplification and resequencing were designed based on *de novo* assembled near-complete SBuMV genome transcripts. At least four successful 5′ RACE clones were sequenced, and according to the obtained sequence alignment onto the *de novo* assembled transcripts from HTS data, the 5′ UTR was determined to be 133 nt long, starting with guanidine (G). The G at the first position of gRNA could indicate that the genome of SBuMV and other completely sequenced marafiviruses that contain G as the first nt [36-40, 48, 73-78] of their gRNA is 5′ capped when compared with other members of the family *Tymoviridae* which had G at the gRNA 5′ end and were capped with m7G [41]. RNAfold web server [79] analysis demonstrated three putative hairpin structures (Fig 4D) that showed similar secondary RNA structure to that of tymoviruses [79, 80]. The 3′ UTR was 109 nt long (without poly-A tail), demonstrating a putative tRNA-like secondary structure as determined by the RNAfold web server (Fig 4C). Furthermore, the complete genome of SBuMV was validated by resequencing several gRNA segments (Fig 2). As revealed by the HTS data mapped onto the assembled transcripts and Sanger-based resequencing, virtually every genomic feature except for the 5′ and 3′ UTRs (which were missing from the initial *de novo* assembled transcripts) showed a plethora of nt substitutions, some of which were nonsynonymous and resulted in aa changes for ORF products in some of the putative genotypes. However, a high heterogeneity of RNA virus populations termed “viral quasispecies” is not an uncommon phenomenon, which has previously raised questions as to what extent a single RNA virus population-derived sequence, representing major allele frequencies, can describe the mutant clouds present in the sample in reality [81]. Determination of the 5′ and 3′ genome termini showed that the full-length SBuMV genome is a monopartite RNA molecule consisting of 6,989 nt (excluding the poly-A tail).

**Fig 4.**
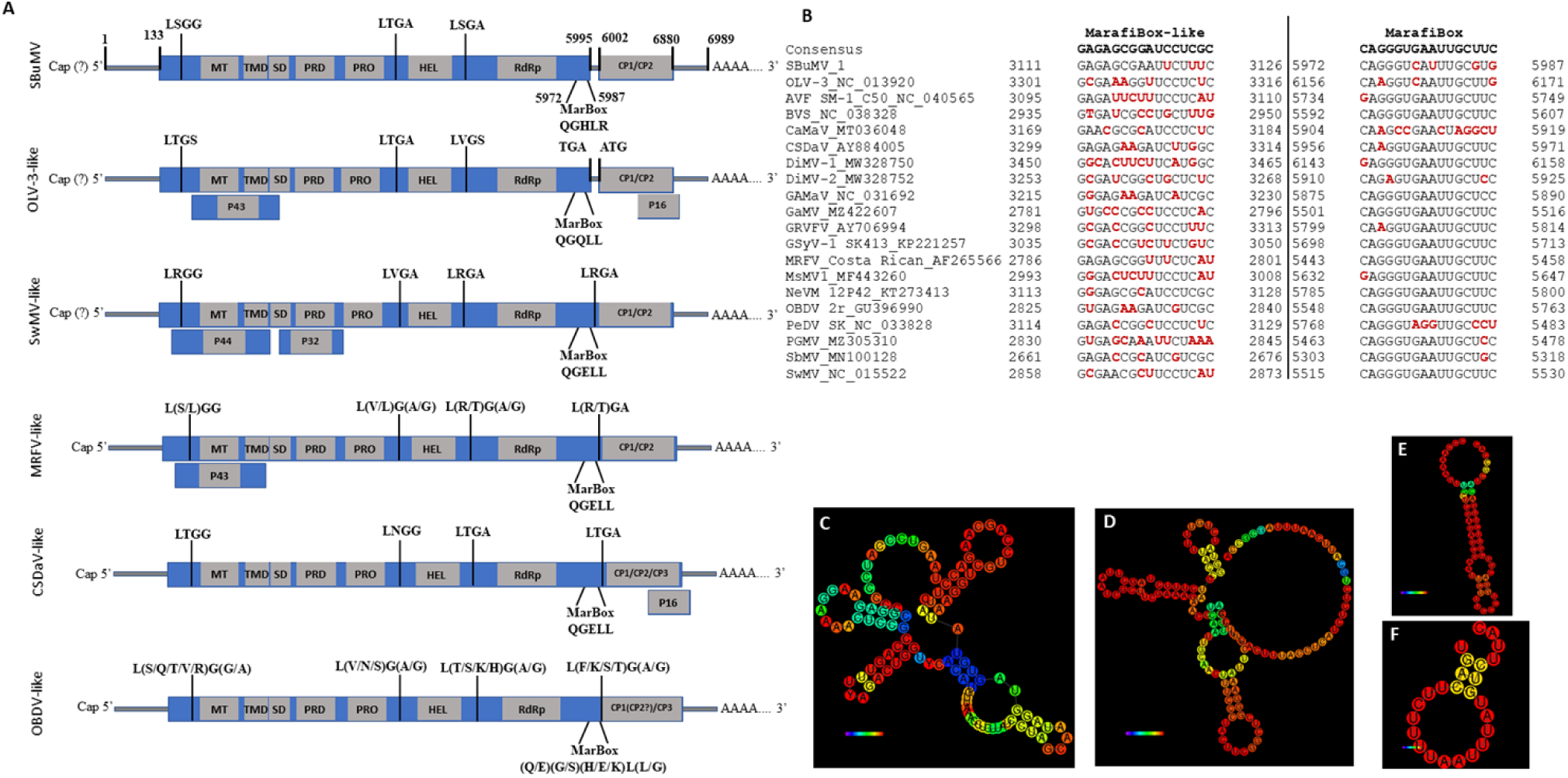
SBuMV genomic features and genome organization. A - Schematic representation of marafivirus genome organizations. MT – methyltransferase; SD - Salyut domain; PRO - protease; HEL - helicase; RdRp - RNA dependent RNA polymerase; MarBox - “marafibox” represented as aa; B - 16 nt “marafibox” and “marafibox-like” sequence alignment; C - predicted RNA structure of SBuMV 3′ UTR; D - predicted RNA structure of SBuMV 5′ UTR; E - predicted secondary RNA structure of SBuMV “marafibox-like” sequence; F – predicted secondary RNA structure of SBuMV “marafibox” Marafiviruses discovered thus far have demonstrated at least six different genome organizations (Fig 4A). Here, we have classified them according to the well-studied marafiviruses: MRFV-like, CSDaV- like, OLV-3-like, OBDV-like, Switchgrass mosaic virus (SwMV)-like, and SBuMV-like. The newly identified SBuMV genome organization resembles that of OBDV, but with a separate ORF encoding a *CP*, which has been observed only in OLV-3 among all *Marafivirus* representatives. The OBDV-like genome organization group consists of six taxonomically recognized members [peach marafivirus D (PeDV), AVF, blackberry virus S (BLVS), NeVM, GAMaV, and Grapevine rupestris vein feathering virus (GRVFV)] and several tentative *Marafivirus* members [MsMV1, DiMV1, Davidia involucrata marafivirus 2 (DiMV2), Camellia-associated marafivirus (CaMaV), Sorghum bicolor marafivirus (SbMV), grapevine-associated marafivirus (GaMV), and Pennisetum glaucum marafivirus (PGMV)]. The MRFV-like genome organization is also characterized by Grapevine Syrah virus 1 (GSyV-1). The CSDaV-like and OLV-3-like genome organization groups only contained their reference members. A new tentative member of the genus *Marafivirus*, *Switchgrass mosaic virus*, possesses a fifth genome organization type (SwMV-like) with the main ORF encoding a polyprotein, as in OBDV-like group, but has two additional ORFs located in close proximity and nested within the polyprotein-encoding ORF (at the 5′ end). This shows that SBuMV genome organization could be viewed as a sixth distinct type of marafivirus genome organization, with SBuMV being the only known marafivirus to possess it. SBuMV ORF1 begins at the 134th nt with methionine and ends at the 5998th nt with a stop codon UAA. The RP was 1,954 aa long, with a calculated molecular mass of 216.5 kDa. RP-encoded domains were determined using the conserved domain database (CDD) [82], suggesting that SBuMV ORF1 contains at least five distinct protein domains: viral MT (pfam01660; 101–382 aa), Salyut domain (cl41199; 550–583 aa), tymovirus endopeptidase (cl05113; 825–920 aa), viral (superfamily 1) RNA HEL (pfam01443; 1014–1245 aa), and RdRp (cl03049; 1661–1854 aa). MSA of SBuMV RP and RP of other officially recognized and selected tentative marafiviruses (20 in total) using the PROMALS3D server [83] allowed us to identify many characteristic domain motifs. Hence, all motifs corresponding to the MT domain (motifs I, II, and III) [84] were identified. Additionally, PRO motifs I (CLL) and II [H(F/Y)], which are conserved between tymo-, marafi-, and macula-viruses and contain a catalytic dyad (C829 and H918), HEL motifs I to VI, and RdRp motifs I to VIII, could be identified according to the OBDV model (S1 File) [37, 38]. The GSyV-1 permuted RdRp motif VI with a conserved viral polymerase aa sequence (GDD) was clearly visible (S1 File) [85]. Two transmembrane domains were identified within the 422–607 aa region and flanked by the putative PRO catalytic sites 422LVGW425 and 604LYGN607. In tymovirus TYMV, an internal sequence (41 aa) of a 140 K protein was previously identified as a chloroplast targeting domain (CTD), similar to SBuMV, which had a C-terminal extension of the MT domain that behaves as an integral membrane protein during infection [86].

ORF2 was in the same reading frame (+2) as ORF1 but was separated by six nt. According to CDD search results, ORF2 encoded a product showing features of a tymovirus-like CP domain (cl03052; 115–288 aa), thus possibly encoding for a CP of SBuMV. The first start codon of ORF2 (minor CP) began at 6,002 nt and the second (major CP) at 6,275 nt; however, both ended at 6,877 nt with a stop codon UAA. The minor CP was 292 aa long and had a predicted molecular mass of 31 kDa (p31), with proline- (29.21%) and serine-rich (15.73%) N-terminal (89 aa), which is similar to OLV-3 CP [37]. This seems to be the largest minor CP variant among not only all marafiviruses, but also among all representatives of *Tymoviridae* [41]. Major CP variant is translated from a second in-frame methionine and the resulting product is 201 aa long, with a predicted molecular mass 21.2 kDa (p21.2). The *CP* gene (ORF2) contains several conserved aa sequences, which are typical for marafivirus CPs – motif I (136PFQW138), motif II (169YRYA174) and motif III (222GGPV225) (S1 File). PFQ conserved aa triplet is present in all of the sequenced viruses belonging to the family *Tymoviridae* [37]. According to the MSA of CP (S1 File), majority of the marafiviruses studied thus far have a second methionine (with the exception of AVF and MsMV1) and a putative PRO cleavage site (except for OLV-3 and SBuMV, which share an MRCA). Interestingly, similar to luteoviruses, SBuMV and OLV-3 have a C-rich region 15 nt after a stop codon, which is a readthrough signal that produces a minor CP [87].

MT (61.5%), PRO (44.14%), RdRp (67.2%), and CP (66.03%) aa sequences of SBuMV shared the highest similarity to the corresponding domains of OLV-3 (YP_003475889.1; [37]). Only HEL shared the highest similarity (66.22%) with the CaMaV (QID59002.1) [76] while having a slightly lower identity (63.46%) with its counterpart from OLV-3. These aa sequence identity values are in accordance with the criteria set for the demarcation of novel species within the genus *Marafivirus* [41]. The SBuMV RP N-terminal contains a 70-aa protein domain with a molecular mass of 7.5 kDa, which is separated from PR by a putative papain-like cysteine PRO cleavage site 61LSGG64, which was also identified in other marafiviruses (S1 File). According to Protein BLAST and CDD results, no similarities were detected with other proteins or their domains, although these might potentially be revealed by employing a more sensitive sequence profile-based search strategy that was not undertaken. RP N-terminal region is also rich in proline (15.87%), serine (14.29%), and threonine (9.59%) residues. PRO cleavage sites between the PRO and HEL domains (929LTGA932) and between the HEL and RdRp domains (1298LSGA1301) were also present.

A “marafibox”-related sequence of SBuMV was located at nt positions 5972–5987, with a putative sgRNA transcription start site of CAAC located at the nt positions 6014–6017. However, CAAU and CAAG were present instead of CAAC in other marafiviruses. SBuMV “marafibox”-related sequences had four changes from the sgRNA 16 nt consensus sequence (5′ CAGGGU<u>C</u>A<u>U</u>UUGC<u>G</u>U<u>G</u> 3′; QGHLR). Differences in this genomic feature have also been reported for other marafiviruses. For example, the PeVD and OLV-3 “marafiboxes” differed from the consensus “marafibox” by six and three nt, respectively (Fig 4B) [37, 73]. The consensus aa sequence encoded by the “marafiboxes” is (Q/E)(A/G/S)(E/Q/K/H)L(L/G/P/R) [76]. Multiple aa sequence alignment of AFV, MsMV1, and DiMV1 “marafiboxes,” where glutamic acid is present instead of a more common glutamine (S1 File), was also consistent with the phylogenetic signals that delineated them into a distinct clade (Fig 2), suggesting that this could already be a feature of their MRCA. The sequence preceding ORF2 formed a secondary stem RNA structure (Fig 4F). Additionally, a sequence similar to the “marafibox” was identified at nt positions 3111–3126 (5′ GAGAGCGAAUUCUUUC 3′). It was located precisely upstream of the HEL domain and is predicted to form a stem-loop RNA secondary structure (Fig 4E). Before it, a slippery sequence motif “XXXYYYZ” (where X represents any three identical nt, Y represents AAA or UUU, and Z represents A, C, or U) was identified, which is a common signal for a programmed -1 ribosomal frameshift (SBuMV: CCCAAAA; 3097–3103 nt) [62, 88]. However, C triplets in such sequences were shown to be the least effective [89].

### Minor and major CP expression in bacterial expression system

SDS-PAGE analysis of p31 and p21.2 revealed that p31 and p31+p21.2 expression levels were reduced compared with that of p21.2 (Fig 5, track T,S). p31, p21.2, and p31+p21.2 sucrose gradient analysis in SDS-PAGE showed that the CPs of all variants were located in 40% and 30% gradient fractions (Fig 5, tracks 3,4).

**Fig 5.**
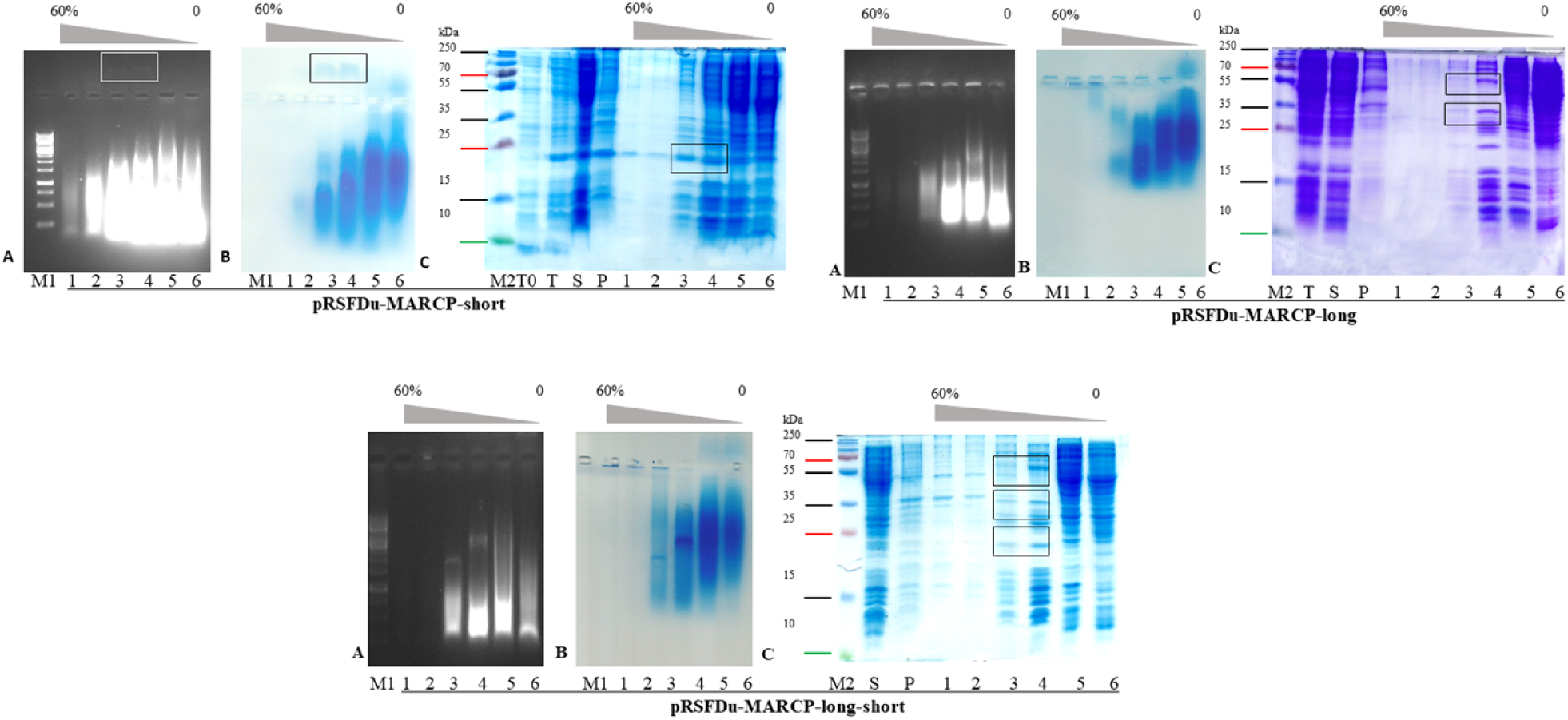
SBuMV native virion and SBuMV VLP purification. A - native agarose gel stained with ethidium bromide; B - native agarose gel stained with Coomassie G250; C - 12.5% SDS-PAGE stained with Coomassie G250 or R250; 1–6 - sucrose gradient fractions starting with 60% (with intervals of 10%) to 20%, overplayed by soluble protein fraction; T0 - total cell lysate before induction; T - total cell lysate after expression of CPs; S - supernatant, which contains the soluble protein fraction after cell disruption by ultrasound; P - pellet in the soluble protein fraction after cell disruption by ultrasound; M1 - GeneRuler 1kb DNA Ladder (Thermo Fisher Scientific, Waltham, MA, USA); M2 - PageRuler™ Plus Prestained Protein Ladder, 10–250 kDa (Thermo Fisher Scientific, Waltham, MA, USA).

Sucrose fractions containing p21.2, p31, and p31+p21.2 were pooled and purified (see virus purification method). Analysis of p21.2, p31, and p31+p21.2 in ethidium bromide stained 0.8% NAG revealed a nucleic acid pattern that migrated towards a negative charge for p21.2, implying that this CP variant has a positive charge (Fig 5, track 1). However, p31 migrated as a negatively charged protein (Fig 5, track 2). After NAG staining with G250, the p21.2 and p31 location signals overlapped with an ethidium bromide-stained band (Fig 5, track 1,2). Notably, p21.2 migrated in SDS-PAGE as a slightly larger protein (Fig. 4, track S) than predicted, whereas p31 showed a migration pattern consistent with a predicted molecular weight of 55 kDa (Fig. 5, track 1). However, both p21.2 and p31 formed additional bands.

Analysis of purified CPs using TEM revealed that p21.2 can readily form homogenous self-assembled icosahedral VLPs with a 30 nm diameter (Fig 6B), resembling typical native marafivirus virions. Furthermore, the analysis of p31-containing fraction revealed only protein aggregates (Fig 6B), while in the case of p31+p21.2, only some partially assembled VLPs were detected (Fig 6B).

**Fig 6.**
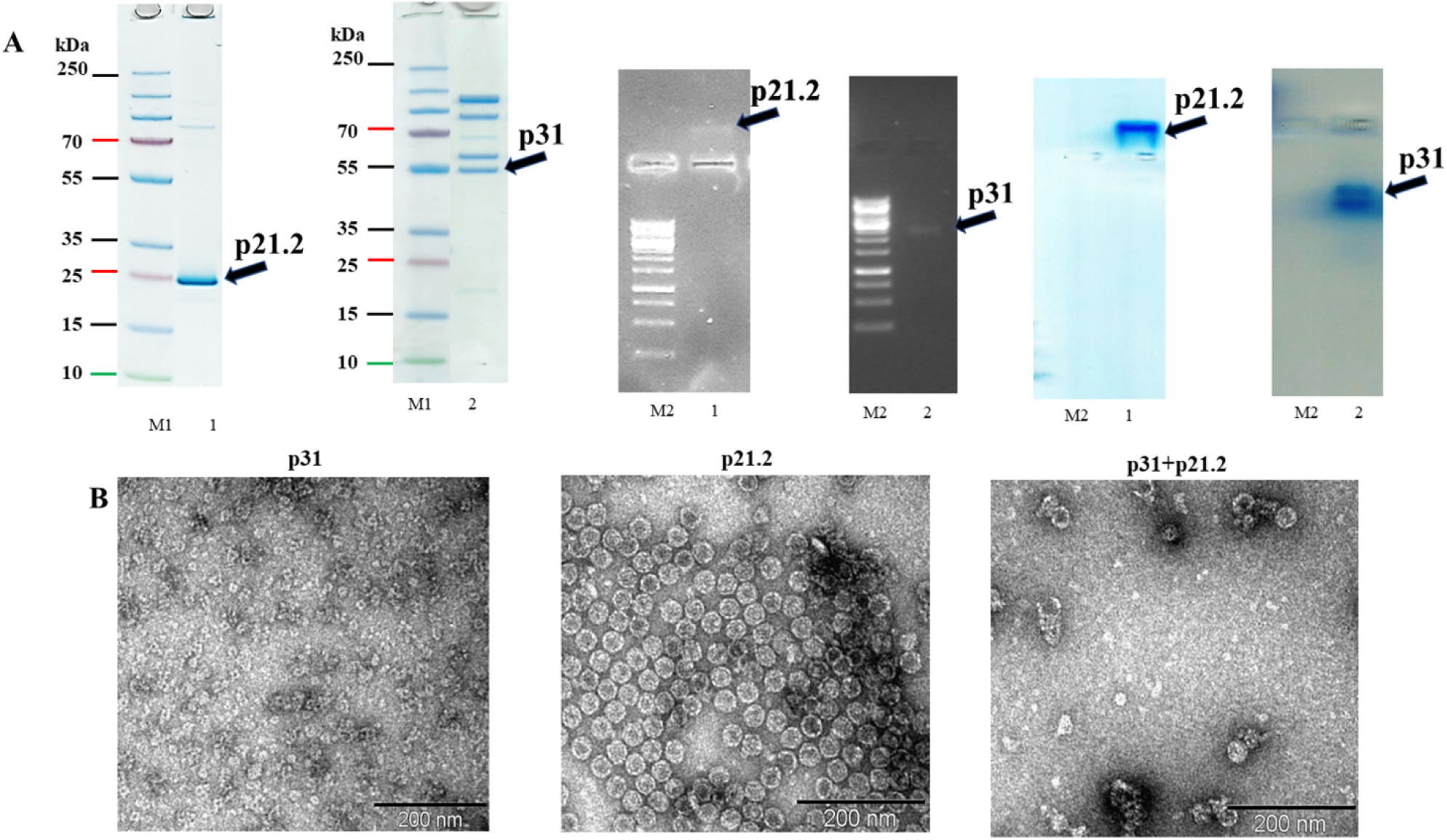
Recombinantly expressed SBuMV CPs analysis. A - SDS-PAGE (stained with G250) and native agarose gel analysis (stained with ethidium bromide and G250); M1 - PageRuler™ Plus Prestained Protein Ladder, 10–250 kDa (Thermo Fisher Scientific, Waltham, MA, USA); M2 - GeneRuler 1 kb DNA Ladder (Thermo Fisher Scientific, Waltham, MA, USA); 1 - purified p21.2; 2 - purified P31; B – Transmission electron microscopy analysis, scale 200 nm.

In case of MRFV, VLPs were obtained by refolding the CP from the inclusion bodies, which could lead to CP proteolysis and possible VLP formation in case of minor CP. Here, SBuMV VLPs self- assembled directly in the bacterial cells, and the majority of the expressed CPs were soluble. The results of this study suggest that the minor CP is not essential for the assembly of seemingly structurally intact viral particles, meaning that it can have other functions.

### SBuMV detection in a follow-up samples

RT-PCR was used to detect the presence of genetic material for SBuMV minor CP-encoding region. All three of the collected samples were found to be SBuMV positive through RT-PCR analysis in 0.8% NAG (Fig 7).

**Fig 7.**
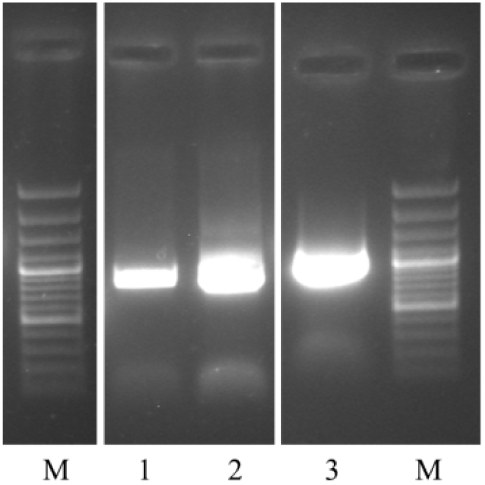
RT-PCR-based SBuMV detection in the follow-up samples. 1–3 - SBT samples collected in 2021; M - GeneRuler 100 bp Plus DNA Ladder (Thermo Fisher Scientific, Waltham, MA, USA).

## Discussion

The discovery of novel viruses provides information that allows for a better understanding of the mechanisms of viral replication, translation, particle assembly, and movement, which are essential for their use as tools for biotechnological applications. Additionally, knowledge of novel possible causative agents of these diseases is invaluable for epidemiology and outbreak containment. Undeniably, the discovery of novel viruses also increases the global understanding of viral diversity, which enables extensive evolutionary studies and further improves the possibilities for conducting comparative genomics-based studies (such as elucidation of conserved sites within viral proteins and assigning a function to novel viral proteins) that is of utmost importance in the omics era [90].

Moreover, the functional annotation of the SBuMV genome was possible directly due to the fact that many other complete or partial marafivirus genomes have been previously sequenced and are publicly available in the biological sequence repositories. With the growing number of novel viruses being identified using HTS, it can be foreseen that HTS will soon become a conventional method for viral phytopathology and will be applied in diverse studies, complementing classical virological approaches that are slowly becoming obsolete without being complemented by relevant sequence data. A comparison of the marafiviruses uncovered thus far shows that despite having some variation within their genome organization, most of their features are similar. The diversity in the genome organization among the marafiviruses could be due to evolution as a result of adaptation to new host plants; however, sequencing, annotation, and validation errors might also play a role in exaggerating the reality [91, 92]. Additionally, recombination between closely related viruses (some of which might not yet be known to science) within the host cells cannot be excluded when trying to deduce the reasons for such a variety in possible genome organizations within a single genus. RNA recombination was found to mediate the rearrangement of viral genes, repair of deleterious mutations, and acquisition of non-self sequences, resulting in ambiguous phylogenetic signals for some viral taxa when the possibilities of recombination were specifically not accounted for. The evidence for recombination not only between closely related viruses but also between distantly related ones and even between the viral and host RNAs suggests that plant viruses unabashedly test the possibility of recombination with any available genetic material [93]. RNA viruses can be regarded as models of efficiency in compressing the maximum amount of information, such as coding and regulatory signals, into the minimum sequence space. They achieve such efficiency by often employing noncanonical translation mechanisms such as overlapping ORFs, some of which can be very short, and arrangement of the ORFs that can additionally regulate their expression. However, small functional ORFs, often lacking conventional initiation sites, can be difficult to detect. Thus, specialized bioinformatics tools are often required to detect the key viral genes [94]. All marafiviruses encode a large RP with five identical functional domains—MT, SD, PRO, HEL, and RdRp—and the documented differences occur only in the presence of an additional ORF at the 5′ or 3′ end of gRNA. For example, MRFV, OLV-3, SwMV, and GsyV-1 genomes have a second overlapping ORF, which shows low sequence identity to the putative movement protein (MP) of tymoviruses and is missing from other marafiviruses [31]. It is a proline-, serine-, and threonine-rich protein, and was shown to be dispensable for the ability to infect maize and leafhopper transmissibility in the MRFV translation mutant experiments. Moreover, unambiguous evidence that the MRFV ORF43 is expressed *in vivo* is lacking [95]. The absence of *MP* gene could be the result of marafivirus phloem limitation [38]. However, in the genomes of CSDaV, OBDV, Grapevine fleck virus (genus *Maculovirus*; GFkV), and SBuMV, a degenerate p43-like ORF interrupted by several stop codons was identified. Therefore, it is reasonable to hypothesize that the truncated MPs of these viruses are remnants of an ancestral MP that had degenerated during evolution as these viruses became restricted to phloem cells [40]. The N-terminal domain of RP is proline-rich that probably overtakes the function of a missing MP, and the MP can also be expressed from a putative non-canonically translated ORF. This has been observed in members of the family *Luteoviridae*, which are confined to the phloem (vascular tissue) potentially due to their specialized phloem-specific MP. These proteins are translated from a single viral mRNA, sgRNA1, via initiation at more than a single AUG codon to express overlapping genes and by ribosomal read-through of a stop codon [94]. To maximize coding capacity, RNA viruses often encode overlapping genes and use unusual translational control mechanisms [94]. The absence of convincing overlapping ORFs in the SBuMV genome could stem from the inability to identify a novel genome architecture/genetic material organization and/or lack of publicly available relevant protein domain homologs. It is possible that some putative small ORFs may be facilitated by the use of non-AUG initiation codons. Under certain circumstances, the near-cognate codons CUG, GUG, ACG, AUU, AUA, UUG, and AUC are able to support a significant level of initiation (typically 2–15% of the initiation levels from an AUG codon in a similar context), with CUG being the most efficient non-AUG initiation codon in many systems [96, 97]. Initiation by non-AUG codons normally requires a strong initiation context but may also be enhanced in less predictable ways by RNA secondary structures [98]. In several plant viruses, a combination of non-AUG and poor-context AUG initiation codons allows the production of three or even four functional proteins from a single transcript [99–101].

Readthrough of the *CP* stop codon of *Luteoviridae* family was used for minor CP variant translation. The readthrough domain in the N-terminal region, which is conserved, is required for aphid transmission; however, the C-terminus, which is variable, appears to enhance aphid transmission efficiency and function in the phloem retention of the virus, thus influencing systemic infection, virus accumulation, and symptom development [87]. Probably, the N-terminal region of the marafivirus minor CP has similar functions; however, this is yet to be properly tested. This mechanism seems to be especially feasible for OLV-3 and SBuMV because they have separate ORFs for RP and CP, and the CP N-terminal is proline-rich compared with that of other marafiviruses. Although there are no typical PRO cleavage sites before a minor CP, an aa sequence similar to the PRO cleavage site LQGH is present in the “marafibox” of SBuMV, which corresponds to the LQGQ in OLV-3 “marafibox” (S1 File). However, the implications of this *in silico* observation are yet to be verified by functional experiments. Perhaps, this could be a result of adaptation to a different transmission vector, for example, from leafhoppers to aphids. In the case of CSDaV, aphids from CSDaV-positive plants were readily virus-positive, but the leafhopper species collected from the same areas were negative for the presence of virus [40]. In addition, Hammond and Ramirez [36] suggested that MRFV p25 (rich in hydrophobic aa and proline) may play a biological role in the transmission and virion packaging of MRFV. This suggestion was based on examples from other viruses, where N- or C-terminal extensions are involved in virus transmission, assembly, replication, and/or spreading [36], and this suggestion has been recently confirmed [46]. Based on current knowledge, the N-terminus of a minor CP variant, when co-expressed with a major variant, could facilitate the packaging of viral RNA [102, 103]. MRFV and AVF minor *CP* genes were transiently expressed in *Nicotiana benthamiana* [48, 104]. Showing in both cases, that VLPs may encapsulate *CP* mRNA and/or host RNA [48, 103].

The major CPs of marafiviruses are translated from sgRNA by a leaky scanning mechanism [43]. The indirect evidence of VLP self-assembly without a minor CP, and the absence of VLP during heterologous expression in a bacterial system could be associated with the fact that major CP is translated from an sgRNA, which holds true for other marafiviruses. The maturation of minor CP from RP is achieved by PRO processing. Marafiviruses (like tymoviruses) contain a PRO cleavage site. Indirect confirmation for this strategy was observed when AVF minor and MRFV minor and major *CP* variants were either individually expressed or co-expressed by agroinfiltration of *N. benthamiana* [48, 103]. The TEM analysis revealed the presence of isometric MRFV and AVF VLPs of ∼30 nm in diameter (similar to SBuMV VLPs), it was unclear whether the observed VLPs were composed of a minor CP variant alone or of both proteins, where the minor variant could have been post-translationally cleaved by PVX- or host-encoded proteases from the major CP variant. Alignment of the marafivirus RP aa sequences revealed that at least three PRO cleavage sites that appear as a consensus motif LXG[G/A] were clearly identifiable in most of them: the first one at the N-terminal region of RP, the second one between PRO and HEL domains, and the third one between the HEL and RdRp domains. For marafiviruses that have a CP fused with the RP, the fourth site can be identified before a minor CP. *In vitro* translation of MRFV gRNA extracted from virions found in rabbit reticulocyte lysates resulted in the synthesis of polypeptides ranging from 15 to 165 kDa in weight. However, no polypeptides corresponding to the CPs could be detected by immunoprecipitation of the translation products using an antiserum [105]. Translation of the OBDV virion RNA in rabbit reticulocyte lysates resulted in the production of protein domains with molecular masses of 227 kDa (RP with CP), 202 kDa (RP), 133 kDa (MT, PRO, HEL), 94 kDa (RdRp with CP), 70 kDa (RdRp), and 24 kDa (CP), as suggested by the predicted protein sequences [38]. Sequence conservation and similarity to other viruses that utilize papain-like proteases as a means of processing polyproteins and defense against protein degradation via the ubiquitin-proteasome system suggests that similar mechanisms are used by marafiviruses [44]. Comparison of the MRFV PRO 3D structure with known PRO 3D structures revealed turnip yellow mosaic virus (TYMV, family *Tymovirus*) PRO as the closest structural homolog, followed by ovarian tumor domain–containing protein 3 (OTUD3) from *Homo sapiens* and OTUD1 from *Saccharomyces cerevisiae*. Additionally, deubiquitinating activity has been shown for BLVS, CSDaV, GsyV1, MRFV, OBDV, and OLV3 [106]. Based on their overall core fold, MRFV PRO and TYMV PRO seem to be more related to a viral OTU specific ubiquitin hydrolase (DUB) than to papain-like proteases [107, 108]. Tymovirus and marafivirus PRO can be classified as OTU-like cysteine proteases [109]. The tymovirus and marafivirus PRO can be classified as OTU-like cysteine proteases [110]. The MRFV cleavage site LVGA between PRO and HEL domains has been verified in the past, suggesting that this particular site in this location is highly likely to be active for other marafiviruses as well (S1 File) [106]. PRO cleavage sites for several marafiviruses, such as PGMV (INGG), SbMV (VNGG), DiMV2 (LTGS), and DiMV1 (LNGS), differ in either the first or last position (S1 File). Different aa in these positions have also been documented in other phytoviruses. TYMV PRO/DUB recognizes the consensus peptide substrate (K/R)LXG(G/A/S) [111]. Other plant viruses, such as representatives of the *Potyviridae* family, use PRO cleavage that resembles a consensus sequence (Y/F/G)xG(A, N, S) [110].

For many ssRNA(+) viruses, viral gRNAs are capped, allowing for efficient translation in eukaryotic cells. Cellular mRNA capping enzymes of the host are located in the nucleus; thus, many viruses that replicate in the cytoplasm encode their own capping enzymes [112]. Multiple aa sequence alignment of the MT domain demonstrated a characteristic consensus sequence in all the analyzed marafivirus sequences (n = 21; S1 File), and CDD analysis revealed an SD domain [82]. This evidence suggests that all marafiviruses may have a cap structure at the 5′ ends of their genomes. This type of MT is strictly associated with supergroup 3 RdRp and superfamily 1 HEL and can be classified as a tymo-like MT [84].

All positive-sense RNA viruses with a genome size of over 6 kb encode a putative RNA HEL, which is thought to be involved in the unwinding of a duplex during the viral RNA replication and, perhaps, translation as well [113]. Seven conserved superfamily 1 HEL motifs (S1 File) can be clearly identified from the corresponding marafivirus aa sequence alignment, thus confirming that their HEL belongs to superfamily 1 [114]. MT, HEL, and RdRp coding domains are easier to identify because of the presence of more evolutionarily conserved motifs compared with the PRO domain. Conservation of viral papain- like proteases is observed almost exclusively around the catalytic cysteine (C) residue. However, identification of a catalytic histidine (H) is difficult in some of these PRO domains, and no other conserved motifs are detectable in such domains [84]. Although marafiviruses have two clearly recognizable PRO motifs, CLL and GHF (S1 File) [37, 38], there are additional small conserved motifs, such as L(W/R), GL, H(F/L), and L(A/C).

These RNA genomes perform the traditional role of being the blueprint for all viral proteins; however, they also contain *cis*-acting RNA elements (REs) that have been shown to impact many essential viral processes, including protein translation, genome replication, and transcription of sgRNAs. For many viruses, REs are usually located in the 5′ and 3′ UTRs and/or internally within inter-cistronic regions. However, one drawback of this approach is that the size and/or location of the non-coding regions can have some limitations. To avoid such limitations, many plus-strand RNA viruses have evolved to position their REs within their coding regions [115].

One potential drawback of having an RE in a coding region is that the same RNA sequence region physically couples two or more distinct activities; therefore, one or both of the corresponding sequence functions may be compromised compared with a hypothetical free-standing localization. Additionally, in some cases, the relative location of REs within the genome of a given virus may not be optimal for their activity; therefore, compensatory measures may be required. One strategy used by many RNA viruses to deal with sub-optimally positioned REs is to eventually reorganize their relative location within the genome via intramolecular long-range RNA-RNA interactions to mediate translational initiation and viral RNA genome replication and form functional secondary RNA structures [115].

The SBuMV genome contains several putative structural elements in both the 5′ and 3′ UTRs as well as in the coding regions. RNA structure predictions suggested that the 3′ UTR could form a tRNA-like structure similar to that of other viruses. For example, the barley stripe mosaic virus (BSMV, genus: *Hordeivirus*, family: *Virgaviridae*) has a poly-A region followed by a 3′ terminal sequence capable of folding into a tRNA-like structure that can be aminoacylated [116]. The gRNA of tymoviruses contains a 3′-UTR secondary RNA structure that is functionally and structurally related to tRNAs. This peculiar structure allows for tRNA-like domain interaction with tRNA-specific proteins, such as RNase P, tRNA nucleotidyl-transferase, aminoacyl-tRNA synthetases, and elongation factors, thus allowing it to play an important biological role in the viral life cycle [117]. The RNA secondary structure prediction of 5′ UTR suggested the presence of three putative hairpin structures (Fig 4D) that also resemble their counterparts from the tymoviruses, indicating a similar function of the UTR in both.

The internal putative structural element—an additional “marafibox” motif at the beginning of an HEL domain in SBuMV—could be an extra signal sequence. A consensus sequence (GnGAnCGnnUCCUCUC) (Fig 4B) was identified for this putative element from the MSA of the corresponding SBuMV regions to its counterparts from other marafiviruses. An extra “marafibox” motif was also identified in the comparative analysis of GSyV-1, which boasted a secondary motif with an 11 base consensus sequence (CUnnCACUCnC), located at a variable distance from the primary “marafibox”. The conserved RNA sequence of the primary “marafibox” motif is generally found to be foldable into a stem structure topped with a UUCA loop. Relative to the overlying protein-coding frame, the second conserved motif is located at variable distances and in a reading frame that is different from the first motif [118]. Al Rwahnih *et al.* [118] proposed that the second RNA sequence motif in the “marafibox” is not conserved, as it is not an aa coding sequence, but rather a sequence encoding RNA structural information, which is also presumed to be function of the first motif. Before “marafibox,” a slippery sequence motif CCCAAAA corresponding to the common XXXYYYZ (X represents any three identical nt, Y represents AAA or UUU, and Z represents A, C, or U) ribosomal frameshift signal can be identified [62, 88]. Recently, an unusual aspect of encephalomyocarditis virus (EMCV; genus: *Cardiovirus*, family: *Picornaviridae*) translation by a previously undetected -1 PRF site in an internal region of the polyprotein was discovered, despite EMCV serving as a model virus for more than 50 years [119]. PRF is *trans*-activated by viral protein 2A. Consequently, the frameshifting efficiency increases from 0 to 70% (one of the highest known in a mammalian system) over the course of infection, temporally regulating the expression levels of the viral structural and enzymatic proteins [62]. However, separate experiments are required to prove the PRF activity due to the C triplets being the least effective in the slippery sequence and the absence of a “spacer” between the slippery sequence and the PRF [89]. The second explanation for the presence of this motif could be that the “marafibox”, similar to the “tymobox”, is a subgenomic promoter [33]. Several other phytoviruses (e.g., members of *Luteoviridae* family) have three sgRNAs [94].

Pseudoknots, secondary structures resulting from an interaction of stems and loops, represent a structurally diverse group of functional regulatory elements. They play several diverse biological roles, such as forming a catalytic core of various ribozymes, self-splicing introns, and telomerase Moreover, they play a critical role in altering gene expression by inducing ribosomal frameshifting in many viruses [120].

To identify the symptoms of a disease potentially caused by SBuMV, SBT orchids have to be independently tested, and the results should be carefully analyzed considering the possible presence of other potential causative agents so that all of Koch’s postulates are fulfilled. Analysis of the SBuMV genome and protein sequences has raised more questions than it has answered. For example, is the 5′ end capped? Does proteolysis occur at the identified cleavage sites? Which is the vector for SBuMV? All these and many more questions would be answered in the near future, thus paving the way for exciting discoveries and novel knowledge on phytoviruses.

## Author contributions

Conceptualization, I.B., N.Z. and A.Z.; experimental work, I.B., V.Z., N.Z., I.K. G.R., R.L. and J.J., writing—original draft preparation, I.B., N.Z. and A.Z.; writing—review and editing, I.B., N.Z., D.S., I.M.B. and A.Z.; supervision, I.B., A.Z.; funding acquisition, A.Z. All authors have read and agreed to the published version of the manuscript. The funders had no role in study design, data collection and analysis, decision to publish, or preparation of the manuscript.

## Acknowledgments

We are grateful and would like to thank PhD. Monta Brīvība, BSc. Līga Bizdēna, MSc. Kaspars Megnis from the Latvian Biomedical Research and Study Centre’s Genome Centre’s Genotyping and sequencing unit for technical and methodological support during the NGS library preparation and sequencing. We would like to also thank PhD. Davids Fridmanis and MSc. Ivars Silamikelis from the Latvian Biomedical Research and Study Centre’s Bioinformatics core facility for continuous support and advice in the NGS data analysis methodology. We would like to thank Editage (www.editage.com) for English language editing.

## Data Availability Statement

The datasets generated and analyzed for this study are stored on the local server of Latvian Biomedical Research and Study Centre’s Bioinformatics core facility and can be available from the corresponding authors on reasonable request. The dataset used for evolutionary relationship analysis (accessions of the selected publicly available *Tymoviridae* entries and the related sequence metadata) can be seen in **S2 Table**. Generated MSA descriptions and the resulting tree features can be seen in the **S3 Table.**

## Supporting information

**S1 Fig. Maximum-likelihood and neighbor-joining trees generated using *Tymoviridae* capsid protein sequence.** Trees were drawn to scale with branch lengths in units of amino acid substitutions per site. In the case of capsid proteins that were encoded as part of the polyprotein, the last 300 aa of the polyprotein sequence containing presumed CP were used (indicated as “y “_300l”st” next to the accession number of a respective sequence). Tree tip labels are in the form of sequence accession number|virus name|strain and are colored based on the genera to which the virus belongs according to the legend. Two asterisks (**) after the virus name indicate that the virus originated from RefSeq and did not yet have a standing in the official virus taxonomy but was included in the analysis based on taxonomy associated with a RefSeq entry. Three asterisks (***) after the virus name indicate that the sequence originated from the Nuccore database and did not yet have a standing in the official virus taxonomy but was included in the analysis based on taxonomy associated with a complete or near- complete genome GenBank entry that was longer than 6000 bases in length. Botrytis virus F, which is a member of the *Gammaflexiviridae* family, genus *Mycoflexivirus*, serves as an outgroup at which the trees are rooted. The black arrow connects the sea buckthorn marafivirus (SBuMV) leaves in both trees.

**S2 Fig. *Tymoviridae* polyprotein aa sequence with maximum-likelihood and neighbor-joining trees.** Trees were drawn to scale with branch lengths in units of amino acid substitutions per site. In the case of polyproteins that encoded capsid proteins as well, the last 300 aa of the sequence were truncated (indicated “y “_w”CP” next to the accession number of a respective sequence. Tree tip labels are in the form of sequence accession number|virus name|strain and are colored based on the genera to which the virus belongs according to the legend. Two asterisks (**) after the virus name indicate that the virus originated from RefSeq and did not yet have a standing in the official virus taxonomy but was included in the analysis on the basis of taxonomy associated with a RefSeq entry. Three asterisks (***) after the virus name indicate that the sequence originated from the Nuccore database and does not yet have a standing in the official virus taxonomy but was included in the analysis on the basis of taxonomy associated with a complete or near-complete genome GenBank entry that was longer than 6000 bases in length. Botrytis virus F, which is a member of the *Gammaflexiviridae* family, genus *Mycoflexivirus*, serves as an outgroup at which the trees are rooted. The black arrow connects the Sea buckthorn marafivirus (SBuMV) leaves in both trees.

**S1 Table. Primers used for SBuMV CPs and genome fragment amplification and sequencing.**

**S2 Table. The dataset used for evolutionary relationship analysis.** Accessions of the selected publicly available *Tymoviridae* entries and the related sequence metadata are listed. Origin column: “E” indicates that the isolate was listed as an exemplar isolate for the species in ICTV virus metadata resource; “ICTV” indicates that the species is recognized in the official taxonomy of *Tymoviridae*, but no accession was present in virus metadata resource; “RefSeq” indicates that the virus does not yet have standing in the official virus taxonomy but was included in the analysis based on taxonomy associated with a RefSeq entry; “Nuccore>6k” indicates that the virus does not yet have a standing in the official virus taxonomy but was included in the analysis based on taxonomy associated with a complete or near-complete genome GenBank entry that was longer than 6000 bases in length.

**S3 Table. Descriptions of the generated MSA and features of the resulting trees.**

**S1 File. Alignment of marafivirus RP and CP with coded protein domain motifs using PROMALS3D.** h - alpha-helix; e - beta-strand; ***bold and uppercase letters*** - conserved aa; *l* - aliphatic (I, V, L); *@* - aromatic (Y, H, W, F); *h* - hydrophobic (W, F, Y, M, L, I, V, A, C, T, H); *o* - alcohol (S, T); *p* - polar residues (D, E, H, K, N, Q, R, S, T); *t* - tiny (A, G, C, S); *s* - small (A, G, C, S, V, N, D, T, P); *b* - bulky residues (E, F, I, K, L, M, Q, R, W, Y); **+** - positively charged (K, R, H); **-** - negatively charged (D, E); charged (D, E, K, R, H); bold letters - conserved aa motifs; CS - protease cleavage site; highlighted in teal - protease catalytic diad aa, C and H; highlighted in yellow - start of PRO domain expressed in *E. coli*; highlighted in turquoise – tested cleavage site in *E. coli* [1]; MTase-Gtase - methyltransferase-guanylyltransferase; highlighted in bright green - marked conservative aa for MTase-Gtase domain; PRO - protease; TMD – transmembrane domain; Hel - helicase, RdRp - RNA dependent RNA polymerase; CP - coat protein; bold and light blue - putative M for major coat protein; highlighted in grey - sea buckthorn marafivirus (SBuMV) isolate BU1buckthorn marafivirus (SBuMV) isolate BU1.

---

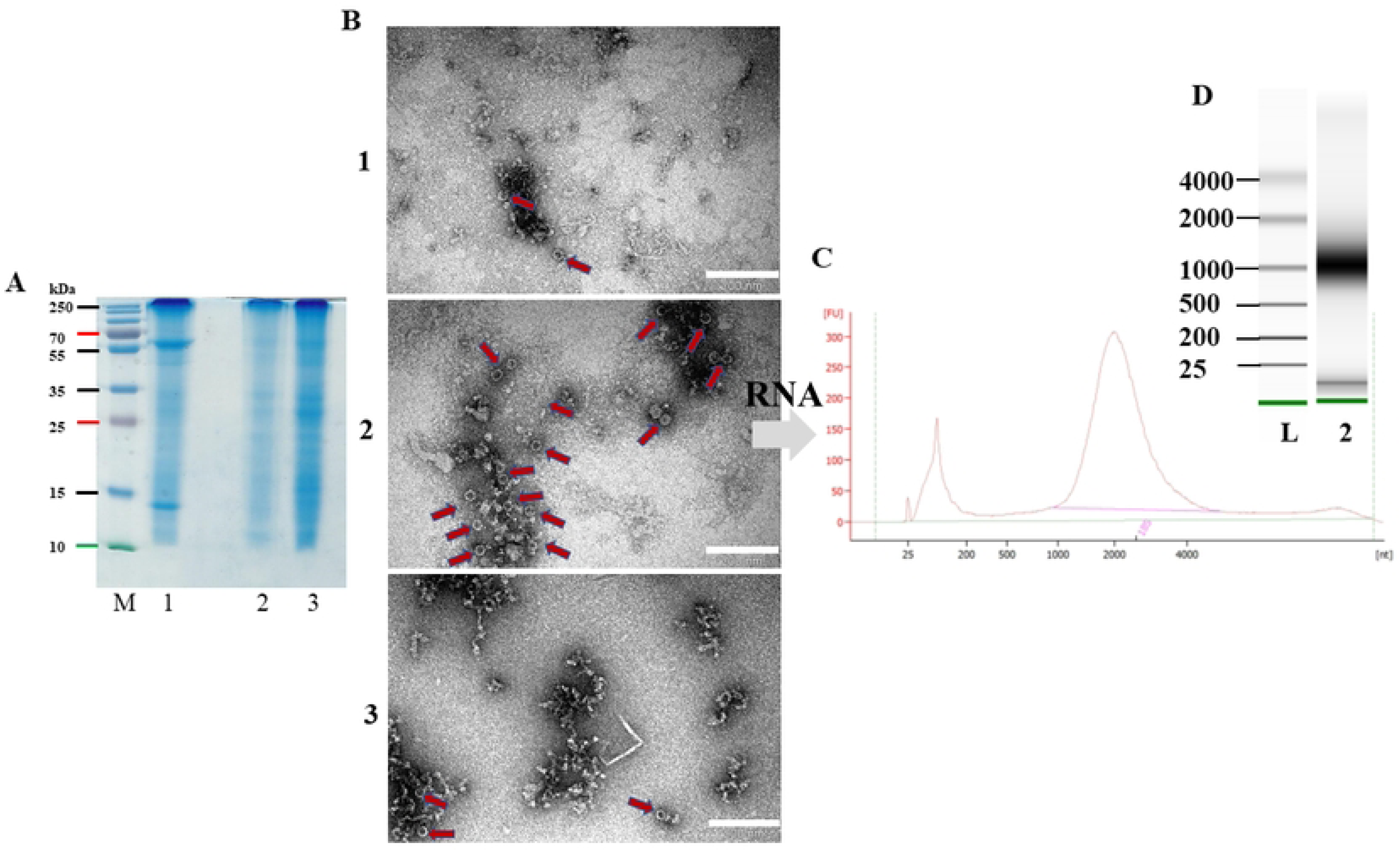

